## Supplementary Information for "Crosslinking-guided geometry of a complete CXC receptor-chemokine complex and the basis of chemokine subfamily selectivity"

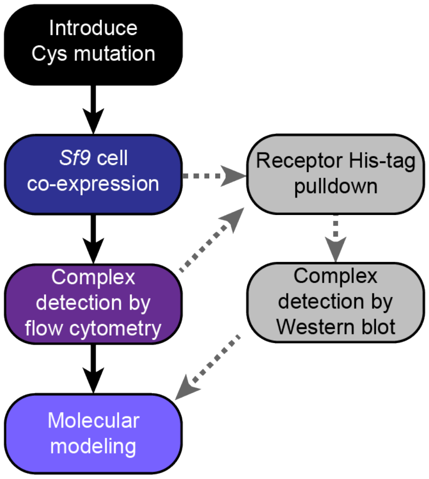


**S1 Fig. Overview of the disulfide-crosslinking-based strategy for determining the geometry of receptor-chemokine complexes in this study.**

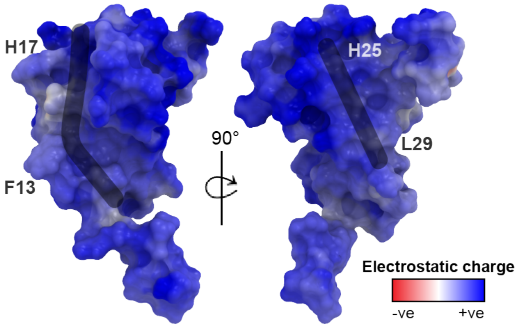


**S2 Fig. The electrostatic surface of CXCL12.** The CXCL12 surface is strongly positively charged (blue). The electrostatic surface was calculated by REBEL in ICM [79]. The black strokes highlight the predicted peptide interaction grooves on the CXCL12 surface: one between the N-/40s loop and another along the β1-strand backbone.

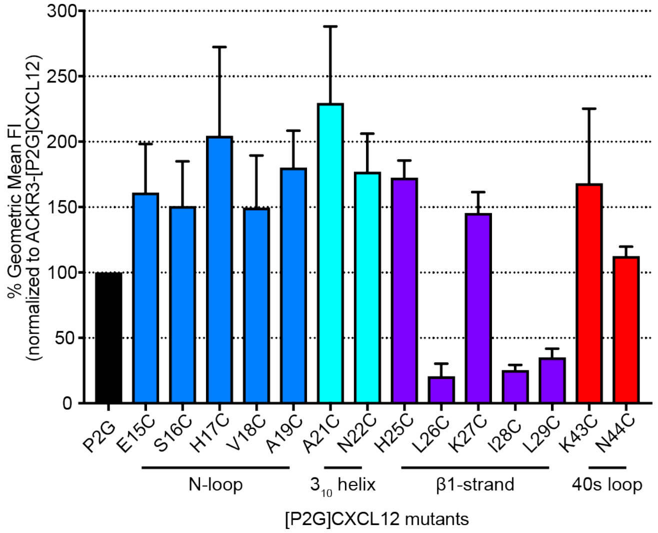


**S3 Fig. Validation of folding of [P2G]CXCL12 Cys mutants by detecting their non-covalent binding to ACKR3.** ACKR3 and [P2G]CXCL12 cysteine mutants were co-expressed in Sf9 cells. Due to the slow off-rate of [P2G]CXCL12 with ACKR3, complexes readily detected on the cell surface are a proxy for mutant chemokine folding. All mutants except L26C, I28C and L29C retain their ability to bind ACKR3. n = 4 independent biological replicates. The mean and s.e.m are reported for each point.

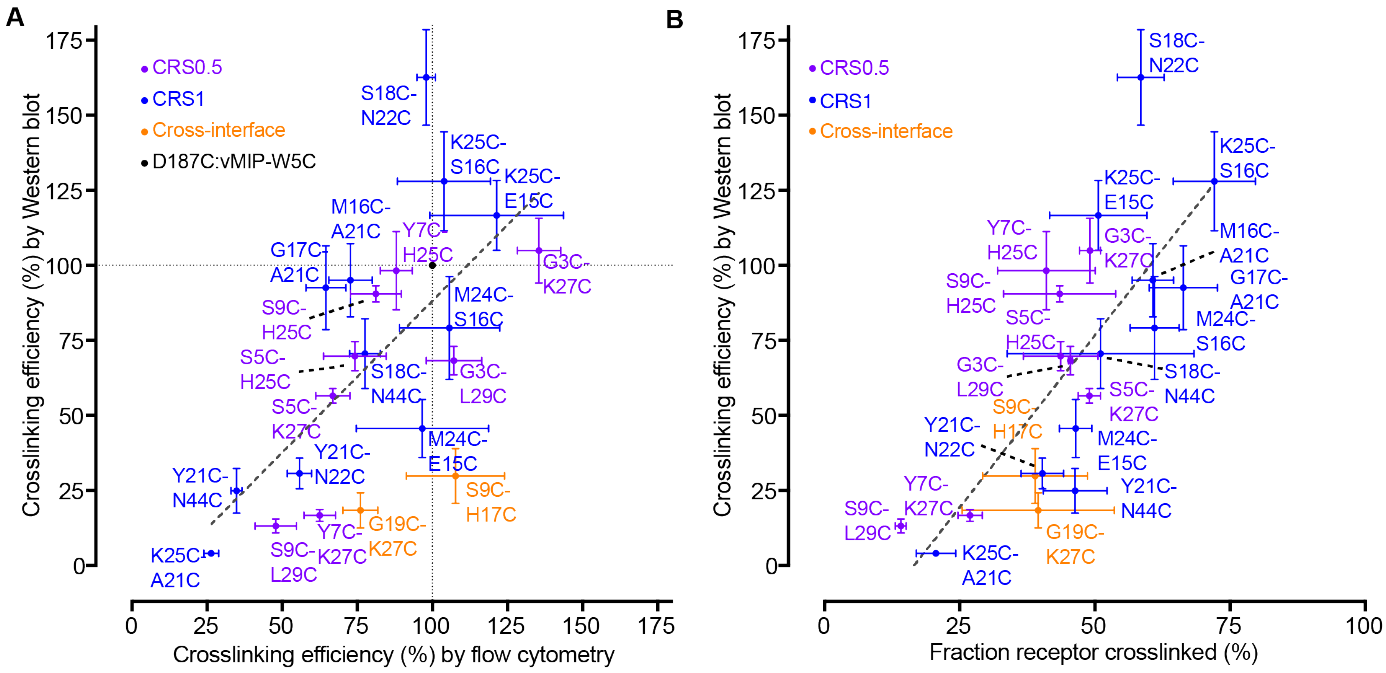


**S4 Fig. Positive relationship between the detection of the chemokine HA tag on the cell surface and in the pulled down crosslinked samples (A), and between the HA intensity and the fraction crosslinked receptor in the pulled down samples (B).** Flow cytometry was used for detection of the chemokine HA tag on the cell surface, while Western blot was used for pulled down protein samples. Data represents mean and s.e.m of three or more independent replicates.

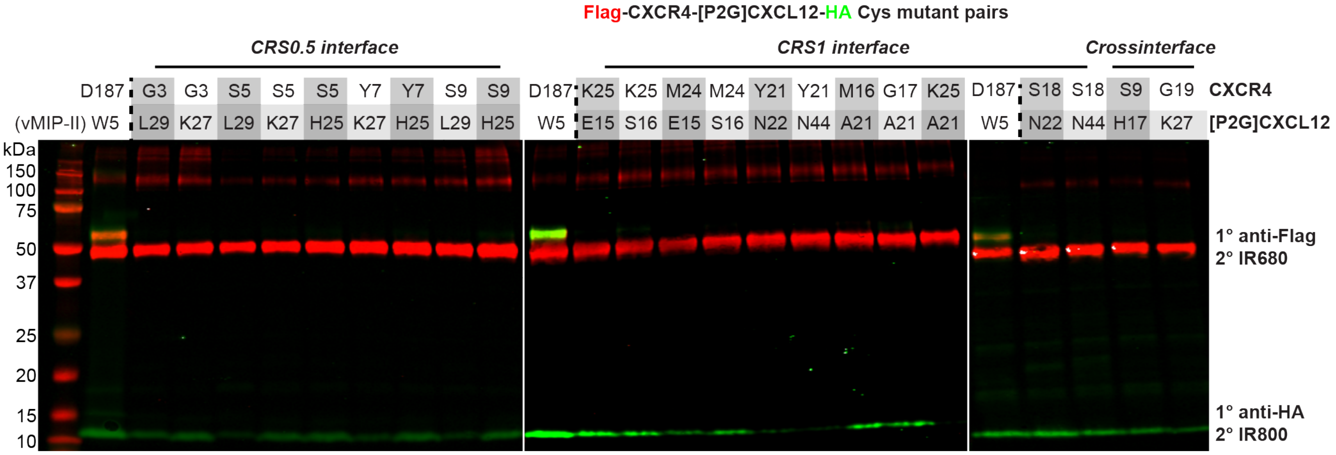


**S5 Fig. The covalent CXCR4-[P2G]CXCL12 complexes dissociate to separate single receptor and chemokine species under reducing conditions.** Western blot of pulled down combinations of CXCR4 and [P2G]CXCL12 cysteine mutants in the presence of 100 mM DTT. The Flag-CXCR4-T4L receptor and [P2G]CXCL12-HA chemokine were detected by LI-COR IRDye® conjugated secondary antibodies on a single blot (emission wavelength of 680 nm and 800 nm, visualized in red and green, respectively). Emitted fluorescence detected at 800 nm and 680 nm from different bands of the Western blot is indicative of a dissociated receptor-chemokine complex. The figure is representative of n = 3 independent replicates.

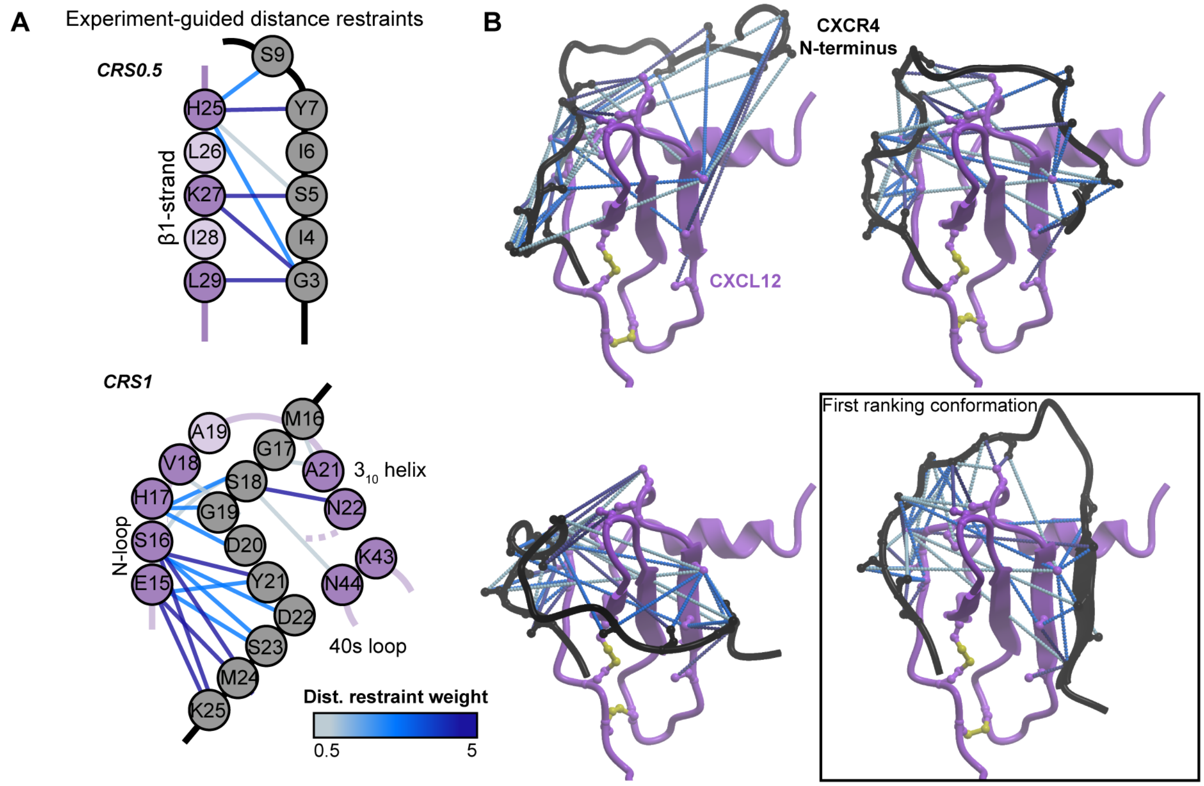


**S6 Fig. The weighted distance restraints imposed between residue Cβ atoms (Cα for glycine) during molecular docking**. (**A**) Graphical representation of the experimentally-derived local distance restraints imposed during the molecular docking simulations. Cross-interface restraints are not shown. Distance restraints are colored by a gradient of blue according to their experimentally-determined strength. (**B**) The distance restraints are mapped onto three randomly selected starting conformations and the top ranked conformation of the receptor N-terminus. Distance restraints are shown in dotted lines, colored by a gradient of blue as in (**A**). The receptor N-terminus and CXCL12 are shown in black and purple ribbon, respectively.

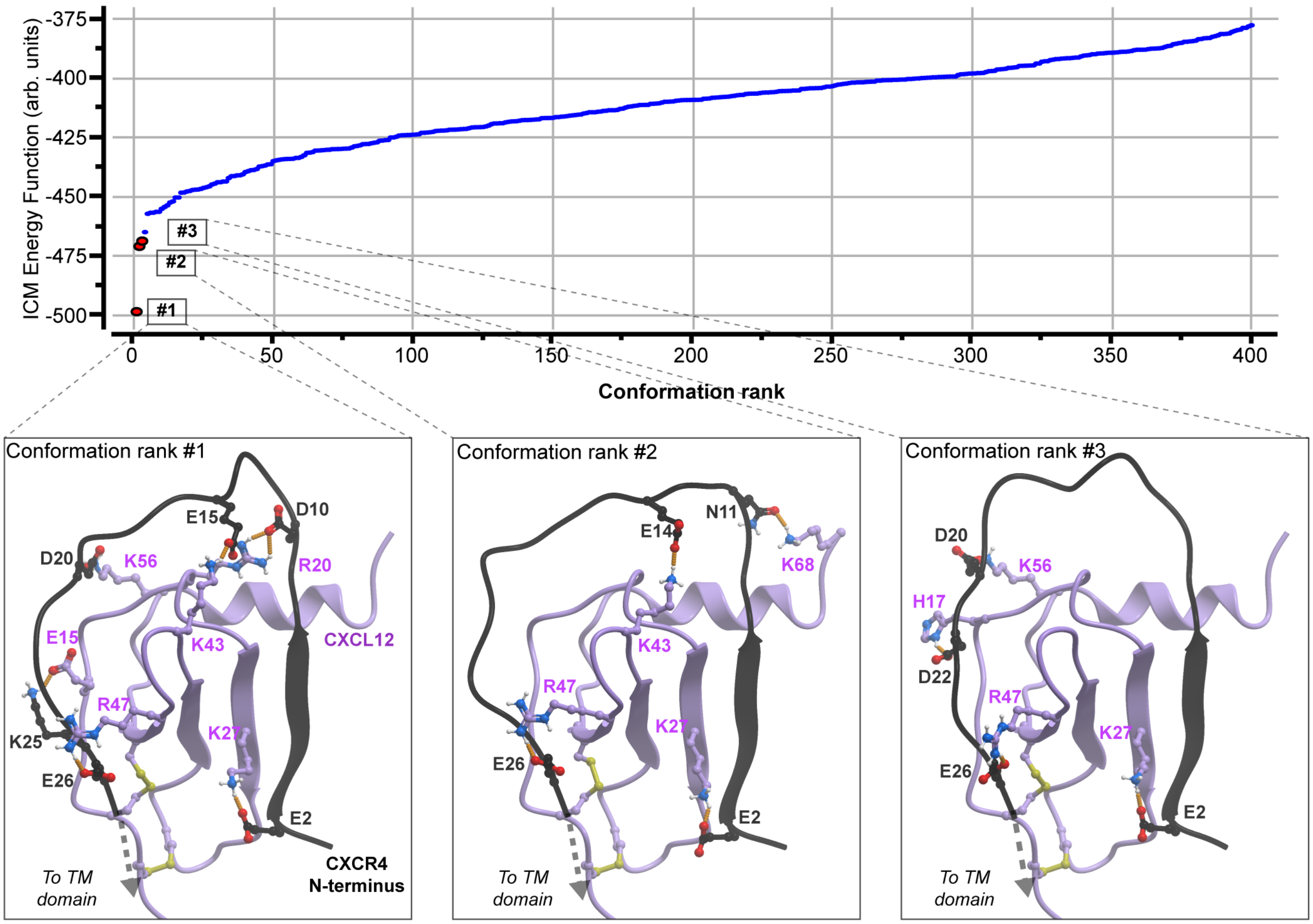


**S7 Fig. The top three ranking conformations of the CXCR4 receptor N-terminus.** The lowest energy conformations are distinct from the other conformations. The conformational stack was sorted by the energy in the system. Polar and charge interactions are shown in orange dotted lines. The receptor and CXCL12 are shown in black and purple ribbon respectively. The TM domain is hidden for clarity.

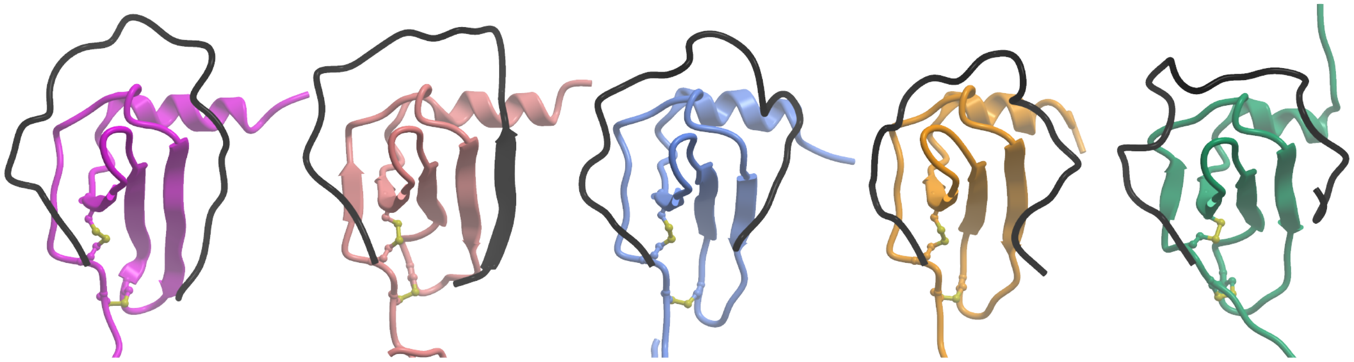


**S8 Fig. The proposed geometry of the receptor N-terminus and the CRS0.5 interface is compatible with various CXCL12 backbone conformations.** The top-ranking conformation from each respective simulation is shown where the receptor N-terminus forms an interface with the CXCL12 β1-strand**.** The CXCL12 conformation PDB 3GV3 from Cluster 2 was selected for full-length complex assembly. The receptor is shown in black, while CXCL12 is colored distinctly in each model.

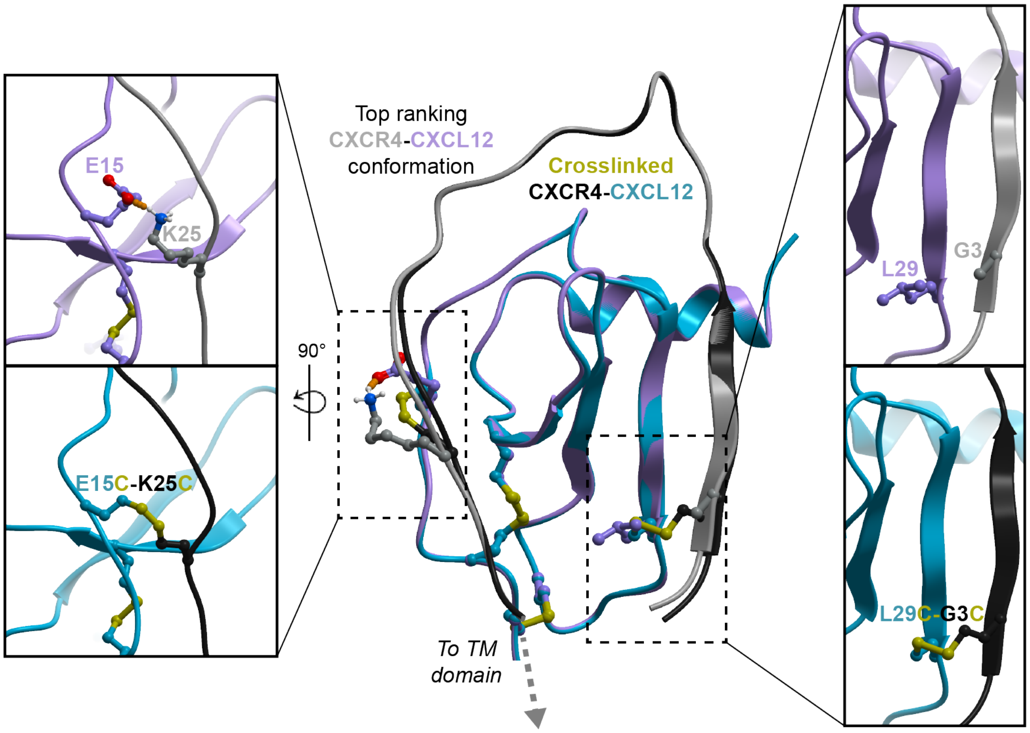


**S9 Fig. Structural context of the crosslinking approach.** Experimental crosslinking observed between the two CXCR4-CXCL12 residues (E15-K25 and L29-G3) can easily be accommodated structurally in our top ranking model with minor changes to the conformation of the receptor N-terminus.

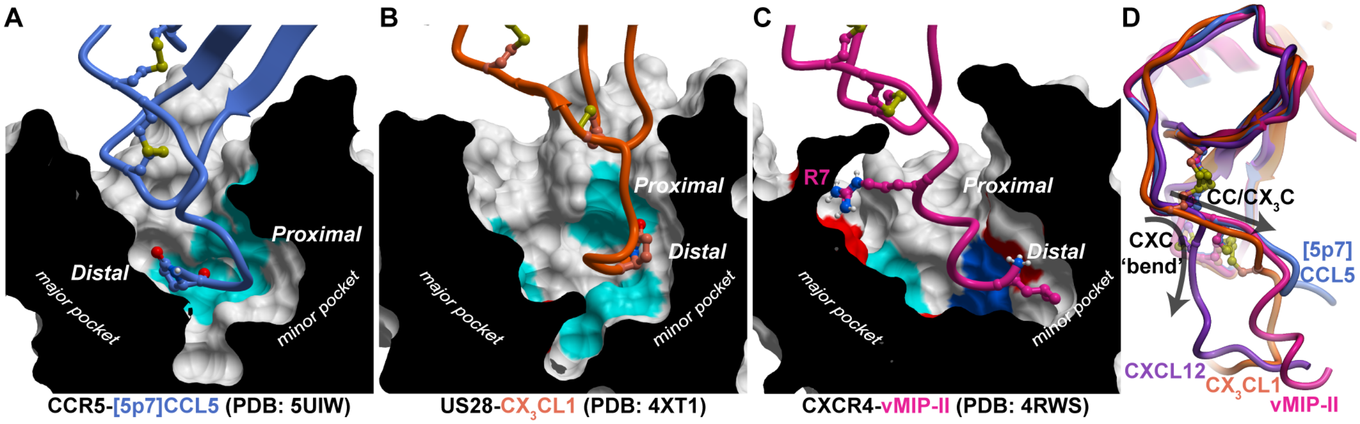


**S10 Fig. Occupancy of the receptor major subpocket by the chemokine proximal N-terminus defines chemokine receptor subfamily selectivity.** (**A**-**C**) Compared to Fig. 4G, the proximal N-terminus of CC and $CX_3C$ chemokines occupy the receptor minor subpocket. vMIP-II is unique among CC chemokines, containing an arginine, conserved in CXC chemokines, allowing it to partially occupy the top of the receptor major subpocket. (**D**) Overlay of the CC and $CX_3C$ chemokines determined in crystal structures, along with CXCL12. CXC chemokines have a pronounced N-terminal bend.

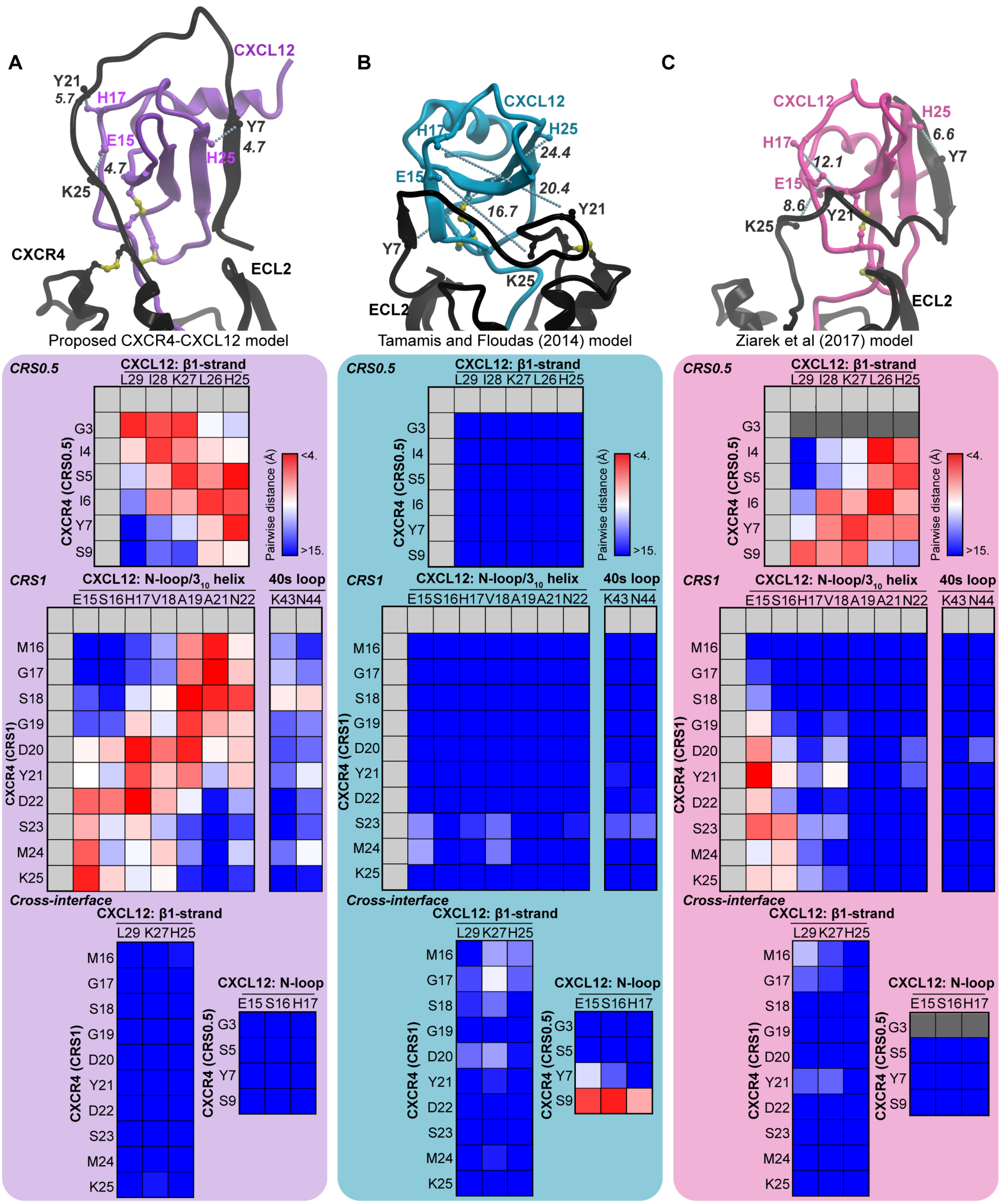


**S11 Fig. Previous published models of CXCR4-CXCL12 complex are incompatible with the crosslinking data.** Receptor-chemokine residue Cβ-Cβ (or Cα for Gly) distances were calculated for three models of the CXCR4-CXCL12 complex and projected onto a heat map for comparison with experimental crosslinking. (**A**) The model generated here; (**B**) the model published by [51]; (**C**) the model published by [25]. We note that the Tamamis and Floudas model was built prior to publication of the CXCR4-vMIP-II crystal structure, and that the Ziarek et al. model was informed by NMR of CXCL12 with an isolated N-terminal peptide of CXCR4. In the Ziarek et al. model, residue G3 was not modelled (dark grey in the heat map). Cβ-Cβ distances between residue pairs (CXCR4 K25-CXCL12 E15, Y21-H17 and Y7-H25) are shown in blue dotted lines, and their distances are given in Ångstroms. The receptor is shown in black, and CXCL12 is colored differently in each model.

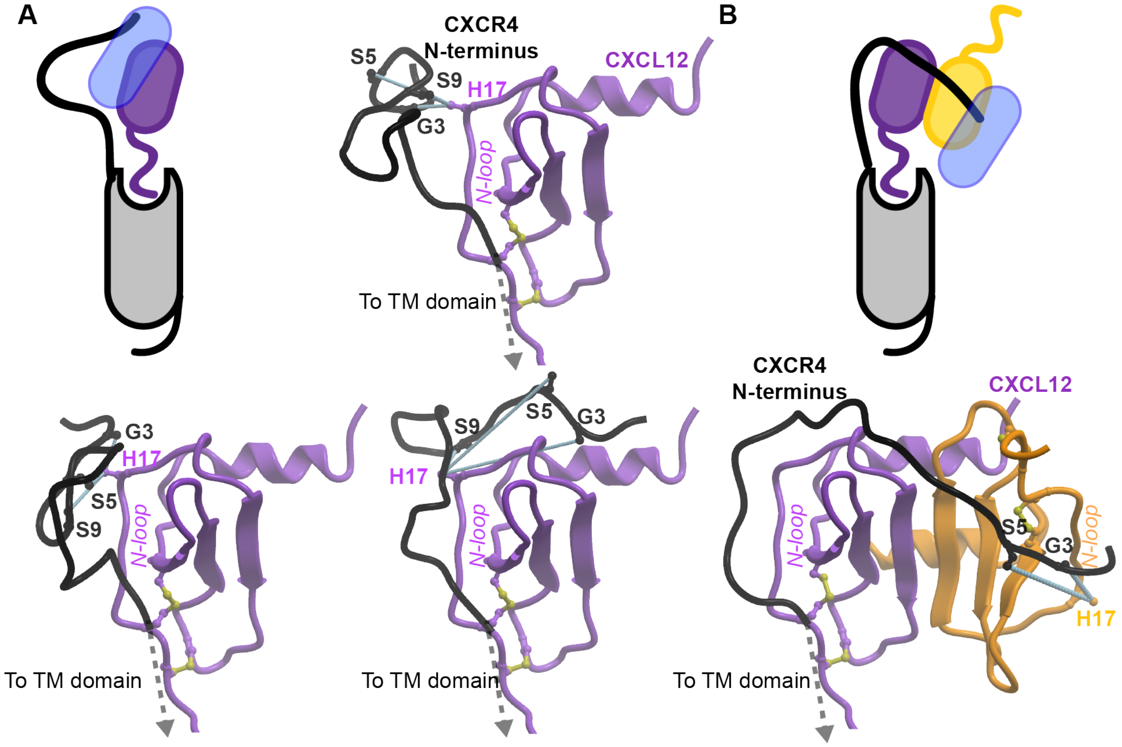


**S12 Fig. Alternative conformations of the CXCR4 N-terminus captured by the docking simulations**. Shown are representative conformations where the distal N-terminus of CXCR4 was found in proximity of the CXCL12 N-loop (**A**) in the context of the CXCL12 monomer or (**B**) in the context of the CXCL12 dimer. In (**b**) case, the distal CXCR4 N-terminus potentially interacts with the N-loop of the CXCL12 dimer partner if fully extended. The receptor and CXCL12 are shown in black and purple, respectively. The second monomer in the CXCL12 dimer is shown in orange. Residue proximities reconciled by these alternative models but not by the best-scoring model are shown as light-blue dotted lines.

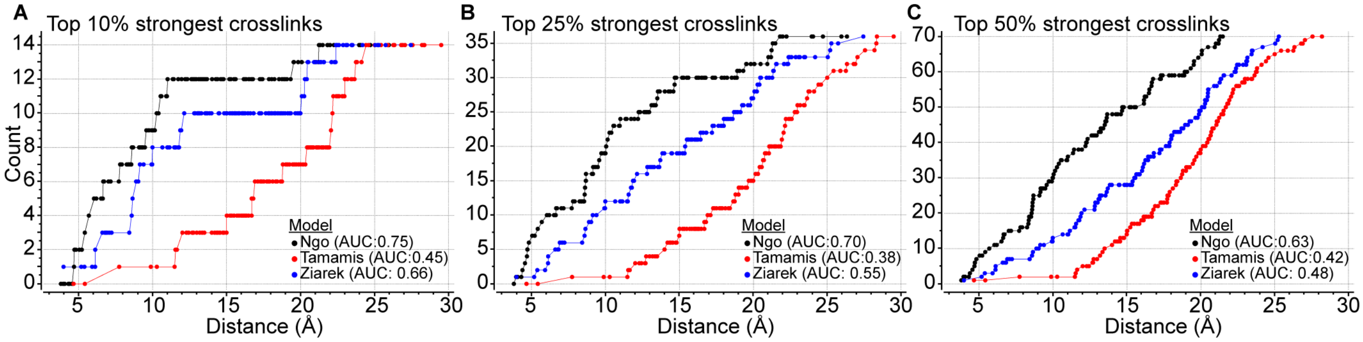


**S13 Fig. Benchmarking comparison of proposed CXCR4-CXCL12 complex geometries.** For each model, pairwise residue Cβ-Cβ distances were ranked and ROC curves were generated based on their ability to recognize: (**A**) the top 10% (14 crosslinks), (**B**) the top 25% (36), or (**C**) the top 50% (72) strongest experimentally determined crosslinks. Model "Ngo" is our proposed model in this study (S1 Data), while "Tamamis" and "Ziarek" are models published previously [25, 51].

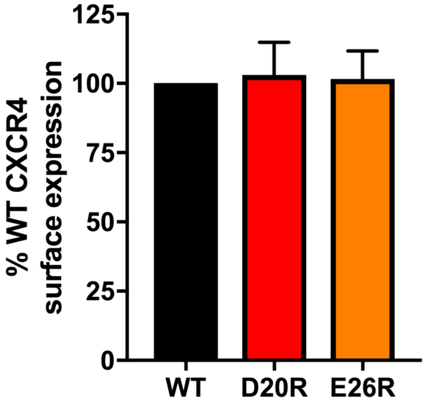


**S14 Fig**. **Surface expression levels of CXCR4 D20R and E26R mutants are comparable to WT CXCR4.** Surface expression was determined by flow cytometry using an APC-conjugated anti-CXCR4 antibody and normalized to WT CXCR4. Data represents mean and s.e.m of n=4 independent replicates.

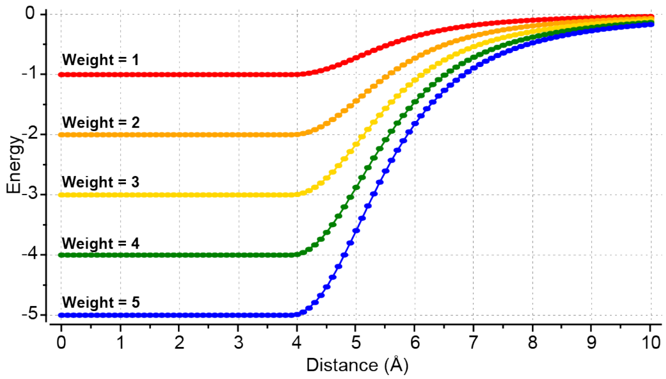


**S15 Fig. The energy function of ICM local distance restraints.** The profile of the restraint penalty features a flat well of varying depth (defined by the restraint weight) at shorter interatomic distances, and increases as these distances increase; however, rather than growing indefinitely, the penalty asymptotically approaches 0 as the interatomic distances continue to increase. This allows the sampling procedure to ignore those restraints that cannot be satisfied concurrently with the majority of other restraints, and in this way, resolve conflicts in the experimental data.

**S1 Table. Strategies for determining receptor-ligand complex geometry.**

| Technique | Mapping of interface residue roles | Full-length receptor? | Mutagenesis artifacts | References |
|---|---|---|---|---|
| NMR chemical shifts | Single-sided | Challenging | N/A | [26, 27, 31] |
| NMR NOEs | Pairwise | Challenging | N/A | [24, 25, 30, 49] |
| Transferred cross saturation NMR | Single-sided | Feasible | N/A | [56] |
| Solid state NMR | Single-sided | Feasible | N/A | [57] |
| Site-directed mutagenesis | Single-sided | Feasible | Possible | [22, 24, 25, 33, 34, 43] |
| Radiolytic footprinting | Single-sided | Feasible | N/A | [34] |
| **Pairwise charge-swap mutagenesis** | **Pairwise** | **Feasible** | **Possible** | **[22]** **This study** |
| **Disulfide crosslinking** | **Pairwise** | **Feasible** | **Possible** | **This study** |

**S2 Table. Physicochemical properties of chemokine CXCL12 based on its crystal structures.**

| Residue | Domain | Flexibility[a] | Mean B-factor (Å$^2$) | Mean res. SASA (±SD)[b] |
|---|---|---|---|---|
| K1 | N-term | 0.99 | 57.30 | 0.87 ± 0.37 |
| P2 | N-term | 0.98 | 59.46 | 0.82 ± 0.42 |
| V3 | N-term | 0.97 | 55.19 | 0.88 ± 0.47 |
| S4 | N-term | 0.91 | 51.38 | 0.89 ± 0.46 |
| L5 | N-term | 0.91 | 40.65 | 0.86 ± 0.36 |
| S6 | N-term | 0.80 | 32.45 | 0.77 ± 0.36 |
| Y7 | N-term | 0.80 | 32.44 | 0.56 ± 0.28 |
| R8 | N-term | 0.72 | 37.91 | 0.78 ± 0.20 |
| C9 | CxC | 0.30 | 28.66 | 0.30 ± 0.09 |
| P10 | CxC | 0.30 | 28.52 | 0.44 ± 0.12 |
| C11 | CxC | 0.21 | 30.13 | 0.08 ± 0.06 |
| R12 | **N-loop** | 0.75 | 40.35 | 0.74 ± 0.15 |
| F13 | **N-loop** | 0.52 | 28.28 | 0.68 ± 0.09 |
| F14 | **N-loop** | 0.52 | 32.16 | 0.40 ± 0.07 |
| E15 | **N-loop** | 0.32 | 32.58 | 0.36 ± 0.09 |
| S16 | **N-loop** | 0.38 | 34.73 | 0.45 ± 0.10 |
| **H17** | **N-loop** | **0.39** | **39.05** | **0.86 ± 0.05** |
| V18 | **N-loop** | 0.31 | 24.98 | 0.10 ± 0.04 |
| A19 | **N-loop** | 0.24 | 25.29 | 0.47 ± 0.05 |
| R20 | **$3_{10}$ helix** | 0.53 | 34.49 | 0.48 ± 0.10 |
| A21 | **$3_{10}$ helix** | 0.22 | 25.92 | 0.67 ± 0.06 |

| | | | | |
|---|---|---|---|---|
| N22 | **$3_{10}$ helix** | 0.34 | 28.12 | 0.29 ± 0.08 |
| V23 | - | 0.32 | 22.49 | 0.08 ± 0.04 |
| K24 | - | 0.42 | 28.00 | 0.51 ± 0.07 |
| H25 | **β1-strand** | 0.43 | 26.26 | 0.42 ± 0.08 |
| **L26** | **β1-strand** | **0.25** | **22.65** | **0.16 ± 0.07** |
| K27 | **β1-strand** | 0.56 | 28.45 | 0.49 ± 0.09 |
| **I28** | **β1-strand** | **0.33** | **22.56** | **0.44 ± 0.06** |
| **L29** | **β1-strand** | **0.34** | **26.63** | **0.22 ± 0.05** |
| N30 | 30s loop | 0.51 | 29.32 | 0.83 ± 0.10 |
| T31 | 30s loop | 0.47 | 29.52 | 0.25 ± 0.14 |
| P32 | 30s loop | 0.69 | 30.98 | 0.71 ± 0.10 |
| N33 | 30s loop | 0.62 | 35.04 | 0.89 ± 0.32 |
| C34 | - | 0.37 | 25.63 | 0.12 ± 0.04 |
| A35 | - | 0.46 | 25.64 | 0.69 ± 0.18 |
| L36 | - | 0.67 | 24.47 | 0.34 ± 0.22 |
| Q37 | β2-strand | 0.61 | 29.17 | 0.16 ± 0.12 |
| I38 | β2-strand | 0.23 | 23.08 | 0.06 ± 0.04 |
| V39 | β2-strand | 0.16 | 22.78 | 0.06 ± 0.02 |
| A40 | β2-strand | 0.15 | 20.41 | 0.00 ± 0.01 |
| R41 | β2-strand | 0.48 | 31.34 | 0.31 ± 0.06 |
| L42 | β2-strand | 0.19 | 23.56 | 0.08 ± 0.03 |
| **K43** | **40s loop** | **0.44** | **30.94** | **0.54 ± 0.06** |
| **N44** | **40s loop** | **0.46** | **35.07** | **0.77 ± 0.12** |

| | | | | |
|---|---|---|---|---|
| **N45** | **40s loop** | **0.32** | **32.29** | **0.48 ± 0.08** |
| **N46** | **40s loop** | **0.33** | **33.22** | **0.49 ± 0.06** |
| R47 | β3-strand | 0.57 | 33.84 | 0.61 ± 0.06 |
| Q48 | β3-strand | 0.32 | 29.79 | 0.46 ± 0.06 |
| V49 | β3-strand | 0.22 | 23.06 | 0.11 ± 0.06 |
| C50 | β3-strand | 0.10 | 21.01 | 0.13 ± 0.03 |
| I51 | β3-strand | 0.31 | 24.57 | 0.00 ± 0.00 |
| D52 | - | 0.21 | 27.51 | 0.16 ± 0.06 |
| P53 | - | 0.23 | 28.06 | 0.31 ± 0.10 |
| K54 | - | 0.69 | 32.70 | 0.78 ± 0.07 |
| L55 | - | 0.38 | 27.28 | 0.16 ± 0.05 |
| K56 | αC | 0.65 | 30.14 | 0.80 ± 0.11 |
| W57 | αC | 0.25 | 24.28 | 0.14 ± 0.03 |
| I58 | αC | 0.34 | 25.82 | 0.02 ± 0.02 |
| Q59 | αC | 0.67 | 36.07 | 0.56 ± 0.12 |
| E60 | αC | 0.64 | 37.28 | 0.44 ± 0.09 |
| Y61 | αC | 0.65 | 30.44 | 0.33 ± 0.11 |
| L62 | αC | 0.62 | 26.57 | 0.34 ± 0.13 |
| E63 | αC | 0.74 | 37.08 | 0.53 ± 0.12 |
| K64 | αC | 0.80 | 39.84 | 0.69 ± 0.13 |
| A65 | αC | 0.78 | 33.77 | 0.60 ± 0.23 |
| L66 | αC | 0.87 | 43.14 | 0.74 ± 0.31 |
| N67 | αC | 0.93 | 57.33 | 0.99 ± 0.44 |

| | | | | |
|---|---|---|---|---|
| K68 | αC | 0.99 | 66.76 | 0.98 ± 0.41 |

[a]Normalized range from 0 (rigid) to 1 (flexible) – median = 0.38

[b]Normalized range from 0 (buried) to 1 (exposed) – median = 41.88

**S3 Table. Conformation clustering of CXCL12 crystal structures.**

| PDB ID | Chain | Resolution (Å) | Length | Cluster |
|---|---|---|---|---|
| 1A15 | A | 2.2 | 67 | 1 |
| **1QG7** | **A*** | **2** | **62** | **1** |
| 1A15 | B | 2.2 | 57 | 2 |
| 1QG7 | B | 2 | 66 | 2 |
| 2NWG | B | 2.07 | 64 | 2 |
| 2J7Z | A | 1.95 | 68 | 2 |
| 2NWG | A | 2.07 | 68 | 2 |
| **4UAI** | **A*** | **1.9** | **68** | **2** |
| 3GV3 | A | 1.6 | 63 | 2 |
| 2J7Z | B | 1.95 | 68 | 2 |
| 4UAI | B | 1.9 | 67 | 2 |
| 4LMQ | D | 2.77 | 55 | 3 |
| 3HP3 | C | 2.2 | 64 | 3 |
| 3HP3 | H | 2.2 | 64 | 3 |
| **3HP3** | **F*** | **2.2** | **65** | **3** |
| 3HP3 | I | 2.2 | 65 | 3 |
| 3HP3 | J | 2.2 | 63 | 3 |
| 3HP3 | D | 2.2 | 64 | 3 |
| 3HP3 | B | 2.2 | 62 | 3 |
| 3HP3 | E | 2.2 | 61 | 3 |
| 3HP3 | G | 2.2 | 65 | 3 |

| | | | | |
|---|---|---|---|---|
| **3HP3** | **A*** | **2.2** | **64** | **4** |
| **4LMQ** | **F*** | **2.77** | **58** | **5** |

Each cluster is highlighted with a different color. *Indicates the CXCL12 crystal structure chain of each cluster used as alternative backbone conformations in S8 Fig.

**S4 Table. Crosslinking efficiencies and distance restraint weights of receptor-chemokine cysteine pairs.**

| CXCR4 mutant | Chemokine domain | [P2G]CXCL12 mutant | Crosslinking efficiency % (flow cytometry)[a] | Crosslinking efficiency % (Western blot) | Fraction receptor crosslinked, % (Western blot) | Distance restraint weight[b] |
|---|---|---|---|---|---|---|
| G3C | **β1-strand** | L29C | 107.2 ± 9.3 | 68.3 ± 4.7 | 45.5 ± 0.4 | 3.18 |
| G3C | **β1-strand** | I28C | 19.8 ± 2.7 | ND | ND | - |
| G3C | **β1-strand** | K27C | 135.5 ± 7.2 | 104.9 ± 10.8 | 49.1 ± 1.9 | 5.00 |
| G3C | **β1-strand** | L26C | 22.0 ± 2.4 | ND | ND | - |
| G3C | **β1-strand** | H25C | 81.7 ± 5.9 | ND | ND | 1.54 |
| I4C | **β1-strand** | L29C | 42.7 ± 6.6 | ND | ND | - |
| I4C | **β1-strand** | I28C | 20.7 ± 3.1 | ND | ND | - |
| I4C | **β1-strand** | K27C | 39.4 ± 2.5 | ND | ND | - |
| I4C | **β1-strand** | L26C | 22.7 ± 6.3 | ND | ND | - |
| I4C | **β1-strand** | H25C | 69.9 ± 6.0 | ND | ND | 0.78 |
| S5C | **β1-strand** | L29C | 33.6 ± 2.7 | N/A | N/A | - |
| S5C | **β1-strand** | I28C | 15.9 ± 2.6 | ND | ND | - |
| S5C | **β1-strand** | K27C | 66.9 ± 5.7 | 56.5 ± 2.4 | 49.0 ± 2.1 | 0.58 |
| S5C | **β1-strand** | L26C | 16.5 ± 4.1 | ND | ND | - |
| S5C | **β1-strand** | H25C | 74.2 ± 10.4 | 69.7 ± 4.9 | 43.7 ± 6.9 | 1.06 |
| I6C | **β1-strand** | L29C | 24.5 ± 2.2 | ND | ND | - |
| I6C | **β1-strand** | I28C | 14.0 ± 2.2 | ND | ND | - |
| I6C | **β1-strand** | K27C | 36.5 ± 7.6 | ND | ND | - |

| | | | | | | |
|---|---|---|---|---|---|---|
| I6C | **β1-strand** | L26C | 16.7 ± 1.9 | ND | ND | - |
| I6C | **β1-strand** | H25C | 36.1 ± 5.8 | ND | ND | - |
| Y7C | **β1-strand** | L29C | 27.2 ± 2.2 | ND | ND | - |
| Y7C | **β1-strand** | I28C | 16.8 ± 2.8 | ND | ND | - |
| Y7C | **β1-strand** | K27C | 62.5 ± 5.3 | 16.7 ± 2.0 | 26.9 ± 2.2 | - |
| Y7C | **β1-strand** | L26C | 19.2 ± 2.9 | ND | ND | - |
| Y7C | **β1-strand** | H25C | 88.0 ± 5.3 | 98.2 ± 13.0 | 41.1 ± 9.0 | 1.94 |
| S9C | **β1-strand** | L29C | 47.9 ± 6.9 | 13.2 ± 2.4 | 14.1 ± 1.0 | - |
| S9C | **β1-strand** | I28C | 18.0 ± 4.2 | ND | ND | - |
| S9C | **β1-strand** | K27C | 64.1 ± 11.8 | ND | ND | - |
| S9C | **β1-strand** | L26C | 15.2 ± 3.2 | ND | ND | - |
| S9C | **β1-strand** | H25C | 81.2 ± 8.5 | 90.5 ± 2.7 | 43.5 ± 10.4 | 1.51 |
| M16C | **N-loop** | E15C | 53.1 ± 10.7 | ND | ND | - |
| M16C | **N-loop** | S16C | 50.4 ± 1.8 | ND | ND | - |
| M16C | **N-loop** | H17C | 61.5 ± 2.5 | ND | ND | - |
| M16C | **N-loop** | V18C | 56.7 ± 3.8 | ND | ND | - |
| M16C | **$3_{10}$ helix** | A19C | 51.9 ± 9.1 | ND | ND | - |
| M16C | **$3_{10}$ helix** | A21C | 72.7 ± 7.2 | 95.1 ± 12.2 | 60.7 ± 3.8 | 0.96 |
| M16C | **$3_{10}$ helix** | N22C | 63.4 ± 3.1 | ND | ND | - |
| M16C | **40s loop** | K43C | 22.6 ± 0.6 | ND | ND | - |
| M16C | **40s loop** | N44C | 47.0 ± 7.4 | ND | ND | - |
| G17C | **N-loop** | E15C | 56.6 ± 4.8 | ND | ND | - |
| G17C | **N-loop** | S16C | 48.0 ± 4.7 | ND | ND | - |
| G17C | **N-loop** | H17C | 65.6 ± 7.8 | ND | ND | 0.50 |

| | | | | | | |
|---|---|---|---|---|---|---|
| G17C | **N-loop** | V18C | 66.2 ± 4.0 | ND | ND | 0.54 |
| G17C | **$3_{10}$ helix** | A19C | 48.9 ± 8.2 | ND | ND | - |
| G17C | **$3_{10}$ helix** | A21C | 64.6 ± 6.7 | 92.5 ± 14.0 | 66.4 ± 6.3 | - |
| G17C | **$3_{10}$ helix** | N22C | 57.4 ± 2.4 | ND | ND | - |
| G17C | **40s loop** | K43C | 21.2 ± 0.5 | ND | ND | - |
| G17C | **40s loop** | N44C | 50.3 ± 6.5 | ND | ND | - |
| S18C | **N-loop** | E15C | 52.9 ± 14.2 | ND | ND | - |
| S18C | **N-loop** | S16C | 79.6 ± 14.4 | ND | ND | 1.40 |
| S18C | **N-loop** | H17C | 83.1 ± 4.8 | ND | ND | 1.63 |
| S18C | **N-loop** | V18C | 39.3 ± 6.1 | ND | ND | - |
| S18C | **$3_{10}$ helix** | A19C | 37.3 ± 3.6 | ND | ND | - |
| S18C | **$3_{10}$ helix** | A21C | 67.9 ± 9.0 | ND | ND | 0.65 |
| S18C | **$3_{10}$ helix** | N22C | 97.9 ± 3.1 | 162.6 ± 16.0 | 58.5 ± 4.3 | 2.58 |
| S18C | **40s loop** | K43C | 34.9 ± 3.9 | ND | ND | - |
| S18C | **40s loop** | N44C | 73.4 ± 5.1 | 70.6 ± 11.7 | 51.1 ± 17.3 | 1.00 |
| G19C | **N-loop** | E15C | 41.8 ± 1.8 | ND | ND | - |
| G19C | **N-loop** | S16C | 66.7 ± 8.4 | ND | ND | 0.57 |
| G19C | **N-loop** | H17C | 79.0 ± 10.2 | ND | ND | 1.36 |
| G19C | **N-loop** | V18C | 71.1 ± 3.3 | ND | ND | 0.85 |
| G19C | **$3_{10}$ helix** | A19C | 27.2 ± 2.2 | ND | ND | - |
| G19C | **$3_{10}$ helix** | A21C | 31.9 ± 2.2 | ND | ND | - |
| G19C | **$3_{10}$ helix** | N22C | 58.8 ± 7.0 | ND | ND | - |
| G19C | **40s loop** | K43C | 22.0 ± 1.9 | ND | ND | - |
| G19C | **40s loop** | N44C | 57.3 ± 13.4 | ND | ND | - |

| | | | | | | |
|---|---|---|---|---|---|---|
| D20C | **N-loop** | E15C | 46.5 ± 4.5 | ND | ND | - |
| D20C | **N-loop** | S16C | 52.7 ± 18.4 | ND | ND | - |
| D20C | **N-loop** | H17C | 44.5 ± 8.1 | ND | ND | - |
| D20C | **N-loop** | V18C | 52.0 ± 0.4 | ND | ND | - |
| D20C | **$3_{10}$ helix** | A19C | 37.0 ± 0.8 | ND | ND | - |
| D20C | **$3_{10}$ helix** | A21C | 28.4 ± 3.6 | ND | ND | - |
| D20C | **$3_{10}$ helix** | N22C | 52.0 ± 2.7 | ND | ND | - |
| D20C | **40s loop** | K43C | 24.9 ± 2.9 | ND | ND | - |
| D20C | **40s loop** | N44C | 36.5 ± 9.5 | ND | ND | - |
| Y21C | **N-loop** | E15C | 83.6 ± 18.9 | ND | ND | 1.66 |
| Y21C | **N-loop** | S16C | 111.4 ± 15.4 | ND | ND | 3.45 |
| Y21C | **N-loop** | H17C | 86.0 ± 9.7 | ND | ND | 1.82 |
| Y21C | **N-loop** | V18C | 41.9 ± 4.6 | ND | ND | - |
| Y21C | **$3_{10}$ helix** | A19C | 34.8 ± 2.1 | ND | ND | - |
| Y21C | **$3_{10}$ helix** | A21C | 41.2 ± 4.3 | ND | ND | - |
| Y21C | **$3_{10}$ helix** | N22C | 55.8 ± 4.1 | 30.7 ± 5.1 | 40.3 ± 3.9 | - |
| Y21C | **40s loop** | K43C | 41.2 ± 4.2 | ND | ND | - |
| Y21C | **40s loop** | N44C | 34.8 ± 1.8 | 24.9 ± 7.5 | 46.4 ± 5.9 | - |
| D22C | **N-loop** | E15C | 33.8 ± 4.7 | ND | ND | - |
| D22C | **N-loop** | S16C | 90.2 ± 18.6 | ND | ND | 2.09 |
| D22C | **N-loop** | H17C | 61.1 ± 6.7 | ND | ND | - |
| D22C | **N-loop** | V18C | 35.9 ± 6.2 | ND | ND | - |
| D22C | **$3_{10}$ helix** | A19C | 28.1 ± 0.8 | ND | ND | - |
| D22C | **$3_{10}$ helix** | A21C | 30.9 ± 4.4 | ND | ND | - |
| D22C | **$3_{10}$ helix** | N22C | 56.8 ± 1.8 | ND | ND | - |

| | | | | | | |
|---|---|---|---|---|---|---|
| D22C | 40s loop | K43C | 37.8 ± 4.4 | ND | ND | - |
| D22C | 40s loop | N44C | 28.8 ± 1.5 | ND | ND | - |
| S23C | N-loop | E15C | 84.9 ± 19.1 | ND | ND | 1.74 |
| S23C | N-loop | S16C | 82.4 ± 7.1 | ND | ND | 1.58 |
| S23C | N-loop | H17C | 67.3 ± 9.7 | ND | ND | 0.61 |
| S23C | N-loop | V18C | 44.4 ± 6.6 | ND | ND | - |
| S23C | $3_{10}$ helix | A19C | 36.8 ± 2.9 | ND | ND | - |
| S23C | $3_{10}$ helix | A21C | 34.1 ± 2.6 | ND | ND | - |
| S23C | $3_{10}$ helix | N22C | 63.5 ± 2.3 | ND | ND | - |
| S23C | 40s loop | K43C | 52.1 ± 4.7 | ND | ND | - |
| S23C | 40s loop | N44C | 44.8 ± 3.0 | ND | ND | - |
| M24C | N-loop | E15C | 96.7 ± 22.0 | 45.6 ± 9.7 | 46.5 ± 3.0 | 2.50 |
| M24C | N-loop | S16C | 105.7 ± 16.7 | 79.2 ± 17.1 | 61.1 ± 4.5 | 3.08 |
| M24C | N-loop | H17C | 72.6 ± 7.1 | ND | ND | 0.95 |
| M24C | N-loop | V18C | 32.4 ± 4.3 | ND | ND | - |
| M24C | $3_{10}$ helix | A19C | 36.1 ± 3.1 | ND | ND | - |
| M24C | $3_{10}$ helix | A21C | 34.8 ± 2.0 | ND | ND | - |
| M24C | $3_{10}$ helix | N22C | 43.0 ± 0.9 | ND | ND | - |
| M24C | 40s loop | K43C | 48.4 ± 4.6 | ND | ND | - |
| M24C | 40s loop | N44C | 38.3 ± 3.2 | ND | ND | - |
| K25C | N-loop | E15C | 121.4 ± 22.2 | 116.7 ± 11.7 | 50.7 ± 9.0 | 4.09 |
| K25C | N-loop | S16C | 103.9 ± 15.5 | 128.0 ± 16.5 | 72.1 ± 7.6 | 2.96 |
| K25C | N-loop | H17C | 50.9 ± 5.4 | ND | ND | - |
| K25C | N-loop | V18C | 28.9 ± 3.8 | ND | ND | - |

| | | | | | | |
|---|---|---|---|---|---|---|
| K25C | **$3_{10}$ helix** | A19C | 18.6 ± 0.9 | ND | ND | - |
| K25C | **$3_{10}$ helix** | A21C | 26.4 ± 2.5 | 4.1 ± 0.7 | 20.6 ± 3.7 | - |
| K25C | **$3_{10}$ helix** | N22C | 27.1 ± 1.2 | ND | ND | - |
| K25C | **40s loop** | K43C | 48.6 ± 3.1 | ND | ND | - |
| K25C | **40s loop** | N44C | 23.2 ± 1.4 | ND | ND | - |
| G3C | **N-loop** | E15C | 31.0 ± 3.1 | ND | ND | - |
| G3C | **N-loop** | S16C | 39.6 ± 7.0 | ND | ND | - |
| G3C | **N-loop** | H17C | 69.3 ± 12.8 | ND | ND | 0.74 |
| S5C | **N-loop** | E15C | 30.2 ± 5.3 | ND | ND | - |
| S5C | **$3_{10}$ helix** | S16C | 45.3 ± 7.0 | ND | ND | - |
| S5C | **$3_{10}$ helix** | H17C | 67.7 ± 11.8 | ND | ND | 0.64 |
| Y7C | **$3_{10}$ helix** | E15C | 38.6 ± 4.8 | ND | ND | - |
| Y7C | **40s loop** | S16C | 56.0 ± 5.7 | ND | ND | - |
| Y7C | **40s loop** | H17C | 92.5 ± 13.0 | ND | ND | 2.23 |
| S9C | **N-loop** | E15C | 47.3 ± 4.9 | ND | ND | - |
| S9C | **N-loop** | S16C | 61.9 ± 8.1 | ND | ND | - |
| S9C | **N-loop** | H17C | 107.7 ± 16.3 | 29.8 ± 9.1 | 39.0 ± 9.7 | 3.21 |
| M16C | **β1-strand** | L29C | 26.5 ± 5.9 | ND | ND | - |
| M16C | **β1-strand** | K27C | 42.5 ± 13.5 | ND | ND | - |
| M16C | **β1-strand** | H25C | 68.0 ± 4.7 | ND | ND | 0.65 |
| G17C | **β1-strand** | L29C | 24.4 ± 7.2 | ND | ND | - |
| G17C | **β1-strand** | K27C | 47.5 ± 15.3 | ND | ND | - |
| G17C | **β1-strand** | H25C | 52.7 ± 16.4 | ND | ND | - |
| S18C | **β1-strand** | L29C | 32.8 ± 1.8 | ND | ND | - |
| S18C | **β1-strand** | K27C | 59.9 ± 10.5 | ND | ND | - |

| | | | | | | |
|---|---|---|---|---|---|---|
| S18C | **β1-strand** | H25C | 52.0 ± 19.3 | ND | ND | - |
| G19C | **β1-strand** | L29C | 37.5 ± 2.0 | ND | ND | - |
| G19C | **β1-strand** | K27C | 76.0 ± 5.9 | 18.4 ± 5.9 | 39.5 ± 14.1 | 1.17 |
| G19C | **β1-strand** | H25C | 47.1 ± 17.7 | ND | ND | - |
| D20C | **β1-strand** | L29C | 26.2 ± 8.1 | ND | ND | - |
| D20C | **β1-strand** | K27C | 41.0 ± 15.8 | ND | ND | - |
| D20C | **β1-strand** | H25C | 62.5 ± 4.8 | ND | ND | - |
| Y21C | **β1-strand** | L29C | 45.1 ± 4.0 | ND | ND | - |
| Y21C | **β1-strand** | K27C | 56.5 ± 11.3 | ND | ND | - |
| Y21C | **β1-strand** | H25C | 81.2 ± 12.6 | ND | ND | 1.50 |
| D22C | **β1-strand** | L29C | 32.8 ± 2.1 | ND | ND | - |
| D22C | **β1-strand** | K27C | 58.7 ± 4.6 | ND | ND | - |
| D22C | **β1-strand** | H25C | 62.7 ± 6.7 | ND | ND | - |
| S23C | **β1-strand** | L29C | 41.0 ± 2.6 | ND | ND | - |
| S23C | **β1-strand** | K27C | 52.2 ± 4.6 | ND | ND | - |
| S23C | **β1-strand** | H25C | 69.0 ± 5.4 | ND | ND | 0.72 |
| M24C | **β1-strand** | L29C | 37.6 ± 3.3 | ND | ND | - |
| M24C | **β1-strand** | K27C | 42.2 ± 4.1 | ND | ND | - |
| M24C | **β1-strand** | H25C | 53.2 ± 2.1 | ND | ND | - |
| K25C | **β1-strand** | L29C | 36.8 ± 5.5 | ND | ND | - |
| K25C | **β1-strand** | K27C | 38.4 ± 3.6 | ND | ND | - |
| K25C | **β1-strand** | H25C | 25.6 ± 1.9 | ND | ND | - |

[a] Geometric mean fluorescence intensities (GMFI) for anti-HA ([P2G]CXCL12) or anti-Flag (CXCR4) staining of complex-expressing *Sf9* cells were normalized to CXCR4(D187C)-vMIP-II(W5C)[6] in the same

experiment. Crosslinking efficiencies by flow cytometry are colored by a gradient of blue to red from low to high efficiency.

[b] For receptor-chemokine pairs that displayed >65% crosslinking efficiencies by flow cytometry, anti-HA ([P2G]CXCL12) GMFI were normalized and transformed to distance restraint weights that ranged from 0.5 to 5.

ND = not determined

N/A = not applicable
